# Differential Gene Expression Analysis and Cytological Evidence Reveal a Sexual Stage of an Amoeba with Multiparental Cellular and Nuclear Fusion

**DOI:** 10.1101/2020.06.23.166678

**Authors:** Yonas I. Tekle, Fang Wang, Alireza Heidari, Alanna Johnson Stewart

## Abstract

Sex is a hallmark of eukaryotes but its evolution in microbial eukaryotes is poorly elucidated. Recent genomic studies revealed microbial eukaryotes possess genetic toolkit necessary for sexual reproduction. However, the mechanism of sexual development in a majority of microbial eukaryotes including amoebozoans is poorly characterized. The major hurdle in studying sex in microbial eukaryotes is lack of observational evidence, primarily due to its cryptic nature. In this study, we used a tractable fusing amoeba, *Cochliopodium,* to investigate sexual development using stage specific Differential Gene Expression (DGE) and cytological analyses. Both DGE and cytological results showed that most of the meiosis and sex-related genes are upregulated in *Cochliopodium* undergoing fusion in laboratory culture. Relative gene ontology (GO) category representations in unfused (single) and fused cells revealed functional skew of the fused transcriptome toward DNA metabolism, nucleus and ligases that are suggestive of a commitment to sexual development. While single cells GO categories were dominated by metabolic pathways and other processes indicative of vegetative phase. Our study provides strong evidence that the fused cells represent a sexual stage in *Cochliopodium.* Our findings have further implications in understanding the evolution and mechanism of inheritance involving multiparents in other eukaryotes with similar reproductive strategy.

## Introduction

Sexual reproduction is ubiquitous and considered to have originated in the last common ancestors of all eukaryotes; however, its evolution still remains a mystery particularly among microbial eukaryotes [1–6]. Sexual reproduction can be defined as a stage in the life cycle involving meiosis – a biological process that reduces the genome complement by one-half (haploid); this is followed by fusion of these haploid cells (gametes), in a process commonly known as fertilization, to form a diploid zygote. This definition is primarily based on observations in macroscopic eukaryotes (e.g. animals and plants) that are usually dimorphic with two distinct sexes. Sexual reproduction in macroscopic eukaryotes is well defined with recognizable cellular and molecular signatures [5, 7]. While some variations of sexual reproduction are generally known [1, 8], the nature or existence of sex in most microbial eukaryotes is poorly documented [1]. Microbial eukaryotes including amoebozoans display diverse quality of life cycles that involve various types of asexual reproduction and sexual stages that are usually cryptic [1, 9, 10].

Recent comparative genomic studies of microbial eukaryotes including amoebozoans demonstrate these microbes not only possess sex genes in their genomes, but also these genes are actively being expressed [11–15]. These discoveries debunked the long-held view that microbial eukaryotes were strictly asexual, and solidified the ancestral nature of sex in eukaryotes. However, sexual development and mechanisms of sex in most of these putative sexual lineages is still not well understood [1, 16]. The main hurdle in understanding sexual development in microbial eukaryotes has been a lack of observed sexual activity, probably as a result of their complex and diverse life cycles [9, 17]. Sources of difficulty in observing sexual phases include unculturability under normal laboratory conditions, poorly characterized life cycles, and in some cases the occurrence of sex during a dormant cyst stage, when the cell becomes condensed and opaque, precluding observation or manipulation [18–20]. We recently described a life cycle of a lens-shaped amoeba (*Cochliopodium*) that undergoes extensive multiple cellular and nuclear fusion during active growth, vegetative, stage [16, 21]. *Cochliopodium* possesses a full complement gene toolkit for sex including those that are exclusive for meiosis [11, 13]. This amoeba was suggested to be a likely useful model organism to circumvent some of the hurdles in studying sexual development in microbial eukaryotes with a similar life cycle [16].

The supergroup Amoebozoa includes lineages with diverse life cycles that may or may not involve observable sexual stages. Few lineages of amoebozoans have been confirmed to engage in sexual reproduction [22–28]. Recent genetic studies demonstrate that all amoebae studied possess or express most of the genes that are recognized as exclusive for sex, albeit scanty physical evidence [11]. The four most common life cycle strategies that are assumed to involve sexual stages in microbial eukaryotes are found in the Amoebozoa. These include those that form sexual cysts [18, 19, 28], amoeboid or ciliated reproductive cells [29], trophozoite (vegetative) fusing cells [16], and those that alternate between sexual and asexual morphs [30]. In this study, we will elucidate the sexual development of one of the reproductive strategies with a tractable amoeba, *Cochliopodium,* using cytology and stage-specific differential gene expression (DGE) analysis.

*Cochliopodium* species grow as single cells with a single nucleus through most their life cycle. However, in sufficiently dense cultures they fuse forming larger plasmodial stages, their nuclei migrate within the plasmodium, come into contact and fuse, forming ployploid nuclei [16, 31, 32]. Subsequently, the merged nuclei undergo division, presumably resulting in a new mix of chromosomes in the individual amoebae produced by division during plasmotomy (cell fission). The molecular aspects of this behavior has not been studied. Similar cellular fusion behavior is also known among other lineages belonging to the three major subclades of Amoebozoa [1, 16]. Cellular fusion and nuclear depolyplodization is also reported in other eukaryotic lineages including some mammalian and cancer cells [33, 34]. This cellular behavior is likely ancestral in eukaryotes. Elucidating the molecular aspects of this interesting process in *Cochliopodium* as model eukaryotic microorganism will give insights how microbes with dramatically different life cycles use unorthodox ways to adapt to, and evolve in, changing environments. In this study, we present robust evidence from both cytology and DGE demonstrating the fused stage in *Cochliopodium* is sexual. Fused amoebae are observed to express meiosis genes and other cellular processes that are consistent with sexual development. Our findings will also have implications in understanding the evolution and mechanism of inheritance involving multiple parents in *Cochliopodium* and other eukaryotes that use a similar reproductive strategy.

## Methods

### Sample collection, Single-cell transcriptome sequencing and assembly

Cultures of *Cochliopodium minus* (syn. *C. pentatrifurcatum* [35]) were grown in plastic petri dishes with bottled natural spring water (Deer Park®; Nestlé Corp. Glendale, CA) and autoclaved grains of rice as a food source. Cultures were maintained until they reached maximum-growth density that we have consistently achieved in our laboratory cultures. Unfused (single) cell samples were collected during the first week of culturing before cellular fusion was at its peak (see [16]). Fused cell samples were collected during the steady state of cellular fusion and fission (8-12 days). Fused and single-cell samples were primarily distinguished by their cell size, since fusion status based on ploidy or number of nuclei is difficult to achieve when working with live cells of *Cochliopodium*. Although this is not an ideal approach, based on our previous experience this method gives a rough starting point to classify cells as fused/single [16]. Individual cells (single or fused) were picked using a platinum wire loop (tip) or mouth pipetting techniques and transferred to a clean glass slide or sterile agar medium to wash off bacteria from the amoeba. Cleaned individual amoeba cells (1-10) were transferred into 0.2-mL PCR tubes and processed for sequencing using Seq® v4 Ultra® Low Input RNA Kit (Takara Bio USA, Mountain View, CA) as described in Wood et al. [13]. We also isolated total RNA from 100 cells, from both single and fused conditions, using NucleoSpin^®^ RNA kit (Macherey-Nagel, Düren, Germany) according to the manufacturer’s protocol. The total RNA was processed for sequencing using the SMART-Seq® kit as above. For sequencing, libraries were prepared from 1 ng of cDNA using the Nextera^®^ XT DNA Library Preparation Kit (Illumina Inc., San Diego, CA USA) according to the manufacturer’s instructions. Libraries were quantified on the Qubit^®^ Fluorometer using the DNA HS Assay. All libraries were sequenced at Yale Center for Genomic Analysis on a HiSeq 2500 in paired-end, high-output mode with 75 bp reads.

FastQC (http://www.bioinformatics.babraham.ac.uk/projects/fastqc/) was used to inspect reads of each sample for quality and length. Illumina adaptor sequences and low quality reads with score below 28 were removed using BBDuk (Joint Genome Institute, U.S. Department of Energy, Walnut Creek, CA USA). The trimming of low quality reads from both ends (“rl” trim mode) is based on the Phred algorithm implemented in BBDuk. Using the same program, we also removed reads shorter than 35 bp after trimming. The remaining reads were assembled *de novo* using rnaSPAdes-version 0.1.1 [36] with default parameters for gene inventory and DGE purposes.

### Differential gene expression (DGE) analysis

To quantify the abundances of transcripts in our samples, a rapid and accurate program Kallisto [37] was used, which takes advantage of pseudoalignment procedure robust to error detection. Since there is currently no reference genome data available for *Cochliopodium*, a comprehensive de-novo transcriptome was assembled from multiple samples of *Cochliopodium minus* [35, 38–40]. This master transcriptome was used for mapping the sequencing reads to generate the counts for each sample. The quantification data was then imported using tximport package in R for downstream DGE analysis. Two DGE tools, DESeq2 [41] and EdgeR [42] that were shown to perform well for smaller number of biological replicates, were initially applied [43]. Based on the results of this initial analysis we designed two additional experimental conditions by selecting samples that showed good clustering and removing bad quality samples (see results). DESeq2 and EdgeR analyses of these experimental conditions showed consistence results with EdgeR showing 91% significantly differentially expressed transcripts overlap with DESeq2 analysis (data not shown). Since DESeq2 has a better moderation in log fold change (FC) for low count transcripts [43], all remaining analyses were performed in DESeq2 packages in R. Fragments per kilobase of transcripts per million base pairs sequenced (FPKM) was used to estimate the level of gene expression. The Benjamini-Hochberg algorithm was used to adjust the resulting p-values for multiple testing with false discovery rate (FDR<0.05) [44]. Genes with an adjusted p-value (padj) of < 0.05 was determined as statistically significant. Up- and down-regulations of the significantly differentially expressed genes (DEGs) were determined using a threshold of 1.5 for log2 FC. Gene clustering of all the DEGs was plotted using function pheatmap in R.

### Gene Inventory and Functional annotation with Blast2GO

To check the expression of potential sexual related genes in different cell conditions, BLAST program was used to search our transcripts with a set of sexual related ortholog groups (OG) obtained from OrthoMCL database (http://www.orthomcl.org/orthomcl/) with E-value cut-off of 10^−15^. The best hits were chosen and checked manually for paralogs using phylogenetic tree. We also used HMMER [45] based homology search as a further confirmation of the genes discovered in our BLAST results. A total of 94 genes were investigated, which included 15 meiosis specific genes, 29 fusion and karyogamy related genes and 50 general sex-related genes based on the previous studies of amoebae [11–13]. The expression of the genes detected in our transcriptome were then further investigated in terms of their fold change and adjusted p-value among different cell conditions using functions in DESeq2 R packages.

To understand the biological activities among different cell conditions, functional annotation of DEGs identified between fused and single samples were performed using Blast2GO [46]. Blast2GO is a bioinformatics tool for high-quality protein functional prediction and provides Gene Ontology (GO) annotation visualization and statistical networks for genetics research. The DEGs that were upregulated in fused and single samples were analyzed using Blast2GO for functional annotation. We first used CloudBlast and InterProScan to search homologs to the DEGs against non-redundant databases from NCBI. GO terms were then retrieved by mapping the hits to the functional information stored in the GO databases. Finally, the obtained GO terms were assigned to the query genes for annotation. InterProScan GOs obtained through motifs/domains were also merged with the results of Blast to the annotations. Enzyme Codes (EC) were also provided when EC number or equivalent GOs are available. Statistical information of the function analysis was checked using the ‘charts’ and ‘graphs’ functions in Blast2GO. Enrichment analysis was performed to check GO terms that were abundant with DEGs upregulated in single or fused cells as test set and all of the DEGs as reference set.

### RNA in situ hybridization (RNA-ISH) of selected sex genes

We used a highly streamlined RNA in situ hybridization, RNAscope (Advanced Cell Diagnostics), method to detect six representative sexual related genes in fixed amoeba cells in all stages. These target genes include meiosis specific (Spo11, Mer3, Dmc1, Pch2), karyogamy (Kem1) and plasmogamy (Bni1). RNAscope involves a series of steps including pretreatments, hybridization, signal amplifications, and detection of target genes. The RNAscope® Multiplex Fluorescent Reagent Kit v2 (Cat. No. 323100; Advanced Cell Diagnostics) was performed using target probes specifically designed for *Cochliopodium* except one gene (Spo11), which was designed using a target sequence from *Acanthamoeba castellanii.* The probes were designed according to the manufacturer’s instructions and include: *20ZZ* probes (Cpe-BNI1-cust targeting 221-1315, Cpe-DMC1-cust targeting 2-987 and Cpe-KEM1-cust targeting 2-1386), *17ZZ* probe Cpe-MER3-cust targeting 1801-2725, *18ZZ* probe Cpe-PCH2-cust targeting 2-965 and *13ZZ* probe Cpe-SPO11-cust 2-676 targeting 2-676 (*A. castellanii*). Actin (*15ZZ* probe Cpe-Actin-cust targeting 104-1076) was used as positive control. Adherent cells representing all stages of amoeba were transferred into a two-well glass chamber slide (Thermo ScientificTM, Nunc Lab-Tek, Rochester, NY) and fixed using either −80°C cooled methanol or 4% formaldehyde (Ladd Research, Williston, VT) for 3–10 min. Fixed cells were dehydrated in a series of ethanol concentrations and then reverse rehydrated before treatment with Hydrogen Peroxide and Protease for permeabilization as stated in the manufacturer’s manual. Hybridization of the probes to the RNA targets was performed by incubation in the HybEZ Oven for 2 hours at 40 ^0^C. After two washes, the slides were processed for standard signal amplification steps for single or multiple channels. Chromogenic detection of probes was performed using OpalTM 520 (FP1487001KT) and OpalTM 570 (FP1488001KT) dyes (Akoya Biosciences) and Hoechst 33358 (1:1500) for DNA. The preparations were examined with a Zeiss LSM 700 Inverted Confocal Microscope (Carl Zeiss MicroImaging, Gottingen, Germany).

## Results

### Single Cell Samples and Transcriptome Data

A total of 20 samples designated as fused (9 replicates) and unfused (11 replicates) were collected (S1 Table). The yield and quality of sequence data differed among our single cells transcriptomes (S1 Table). Generally fused or large cells yielded more sequence data than smaller cells. Since transcriptome sequencing from an individual (one) cell did not generate enough data, likely due to small starting RNA material, we increased the number of cells to 5-10 and collected transcriptome data from these cells using the same approach. We also collected transcriptome data obtained using a regular RNA extraction from 100 cells for both unfused (single) and fused cells (S1 Table). The designations of single and fused state of amoeba cells were primarily made on the basis of amoebae size. However, this was used as a guide rather than a predictor of an amoeba stage. *Cochliopodium* cells show a range of sizes during their life cycle (see [16]). The nucleus is not always easy to visualize due to opaqueness of the cytoplasm and cell inclusions that obscure nuclear observation in live cells. These problems make it difficult in identifying single or fused cells based on size and number of nuclei. Given these challenges we made an effort to select cells based on our observation in previous publications [21, 31, 32] as a general guide to discriminate between single (~20-30 μm) and fused (> 50 μm) amoeba cells.

### Differential Gene Expression (DGE) analysis

PCA (principal component analysis) plot functions in DESeq2 were used to visualize the overall effect of experimental and batch effects and check for possible sample outliers. Two samples that failed the quality check were removed and a plot of the remaining 18 samples showed good clustering for the most part. However, some samples either formed a separate cluster of their own or were mixed with another group (S1 Fig). Taking into account the difficulties in distinguishing cell conditions during our sampling, we applied a strict clustering pattern to select four of the single samples (YT14-YT17) and four of the fused samples (YT26-YT29) that showed good clustering for two experimental conditions (see S1 Table). This two-sample condition analysis, each comprised of 4 replicates, showed good clustering with PC1 representing 76% of variance (Fig 1). Further inspection of the clustering of the 18 samples showed a possibility of an intermediate group consisting of 3 samples (see below).

**Figure 1.**
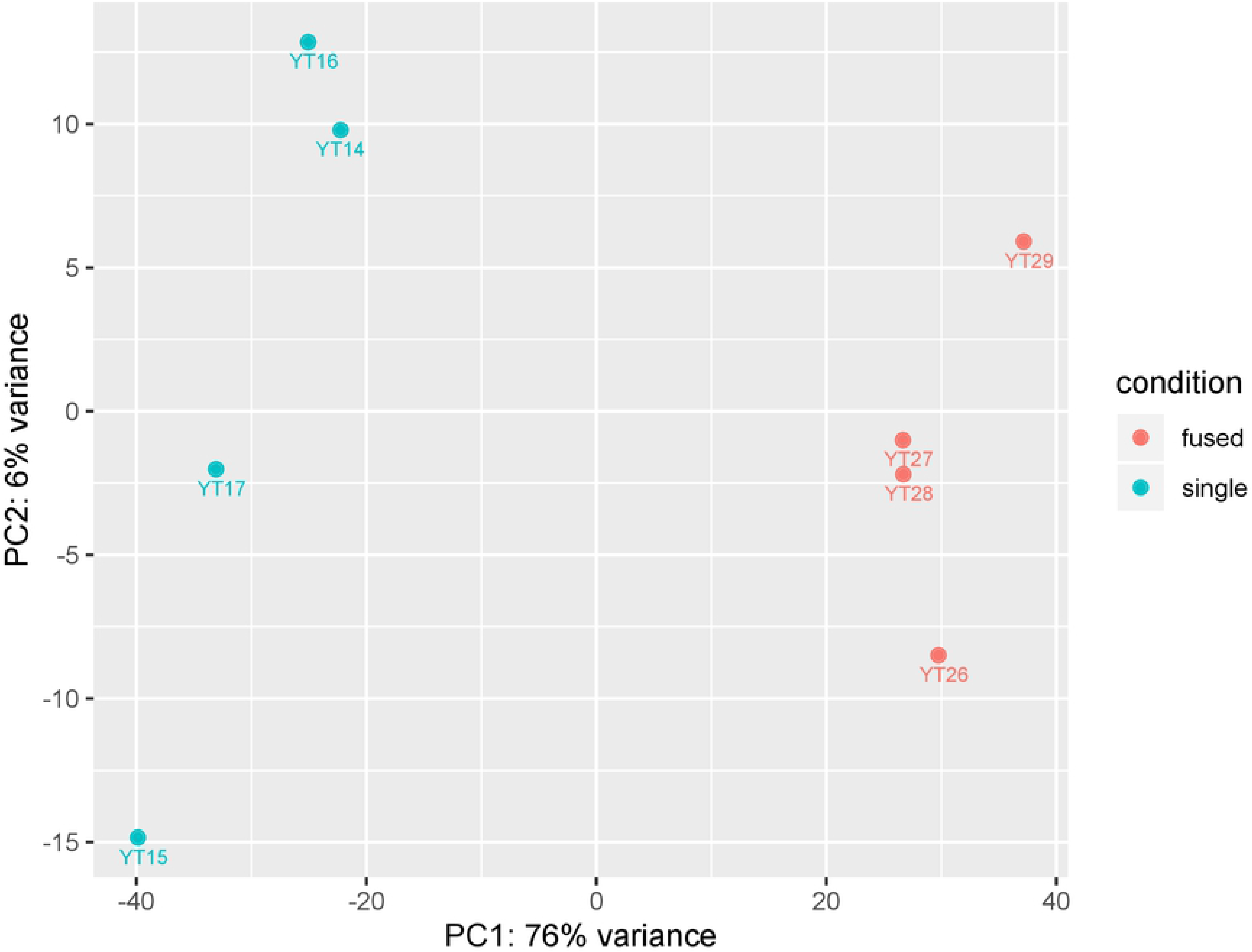
PCA plot of a two -condition experiment (single vs. fused cell) each represented with 4 replicates. X- and Y-axes represent the first and second principal components that explain most of the differences in the data. The percentage of total variance for each principal component is printed in the axis labels.

MA-plot showed that the majority of the transcripts between two-conditions have similar expression values (Fig 2). In total, 881 differentially expressed genes (DEGs) were identified between single and fused conditions in our DESeq2 analysis. Among them, 521 DEGs were tested as upregulated in single condition samples and 360 DEGs tested as upregulated in fused condition samples. The clustered heatmap of all DEGs that were identified between the single and fused conditions was generated using the transformed normalized counts from DESeq2 R package and showed a correlative pattern of upregulation (downregulation) between the two conditions across all samples analyzed (Fig 3).

**Figure 2.**
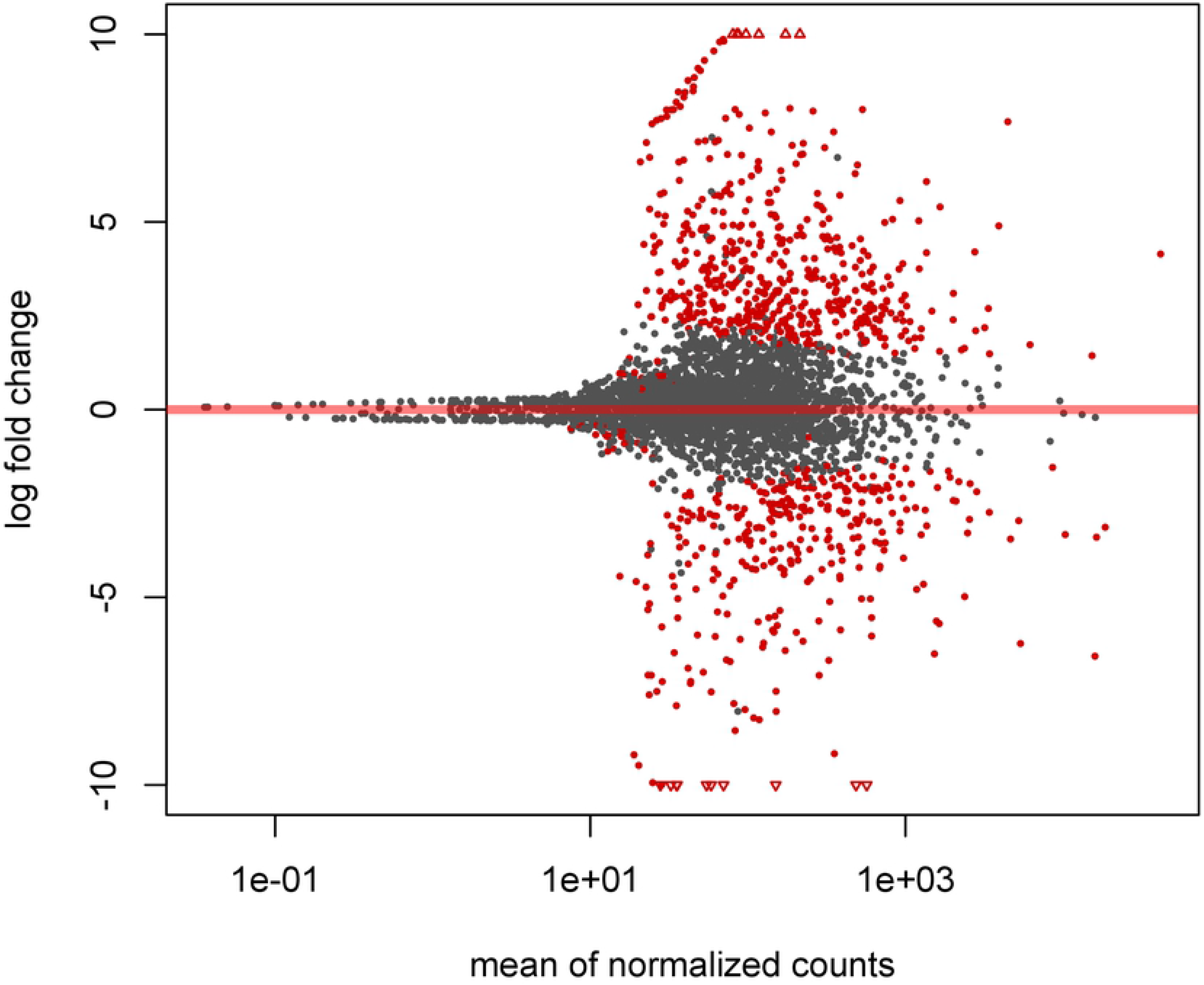
MA plot of the two experimental conditions (single vs. fused) in terms of log2 Fold change and normalized counts. The log2 FC in the y-axis were shrunken to remove the noise from low count genes. The average of the counts normalized by size factor generated in DESeq2 is shown on the x-axis. Points with adjusted p value less than 0.05 are shown as red colors, which is our criterion for DEGs identification. Points that fell out of the window are plotted as open triangles.

**Figure 3.**
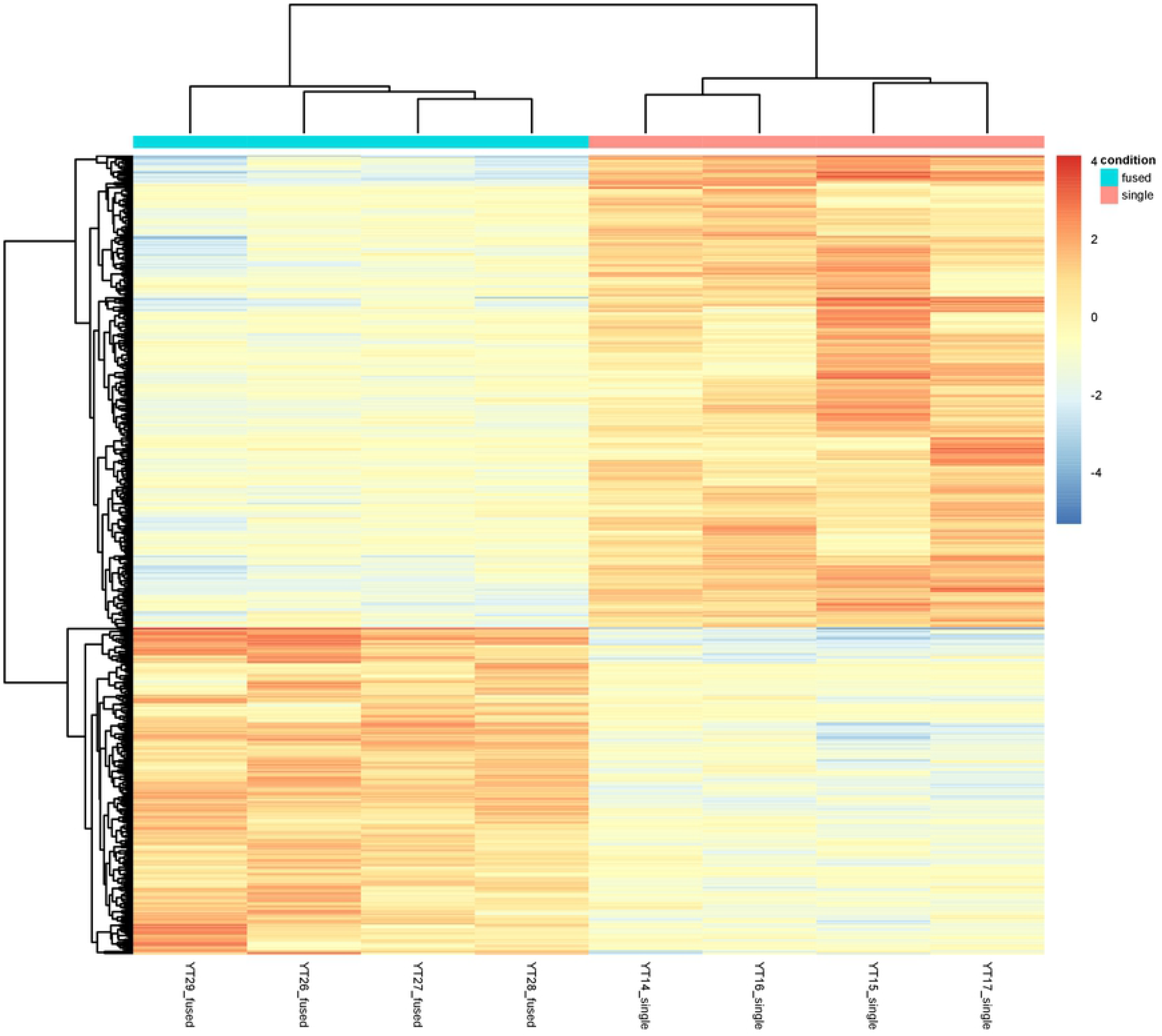
Clustered heatmap of all (881) DEGs identified between single and fused samples. The color scale from red (highly expressed) to blue (low expression) represents the transformed, normalized counts from variance stabilizing transformation.

A closer examination of the expression data structure from our initial analyses revealed a potentially third data cluster (YT24, YT25, and YT30) between single and fused sample conditions (S1 Table, S2 Fig). This cluster likely represents an intermediate condition. This observation is consistent with our previous report on the life cycle of *Cochliopodium* that undergoes a steady state of fusion and fission when growing in high density [16]. Our samples were collected during the active fusion/fission stage and hence some samples might represent intermediate stages despite their cell sizes. To examine the potential role of the intermediate stage in the life cycle of *Cochliopodium*, DGE analysis was performed for a three-conditions experiment (single, intermediate and fused) each represented with three replicates (S2 Fig). In this analysis, a total of 738 DEGs were identified among the three conditions as shown in the Venn diagram of S3 Figure. Among all the DEGs, three separable groups with 113, 54, and 275 DEGs, respectively, were unique for each of the 2 comparisons (see S3 Fig). In specific, there were 287 transcripts upregulated and 229 transcripts down regulated between single and fused stage, while 152 transcripts and 141 transcripts were up- and down-regulated in the single versus intermediate stage (S3 Fig). Comparison between intermediate and fused stage revealed a lower number of DEGs than the above two comparisons, with 131 and 95 transcripts up- and down-regulated, respectively, which indicates the intermediate stage shared more similarity with the fused stage, which is also illustrated in the clustered heatmap of the 738 DEGs (S4 Fig). It can also be said that the intermediate samples exhibited an amalgamation of expression patterns between the single and fused conditions serving as a transitional stage. A set of DEGs are experiencing a gradual increase or decrease in their expression level between single and fused states (S4 Fig).

### Expression of meiosis and sexual related genes

We examined the expression of the genes previously determined to be involved in sexual development of *Cochliopodium* and other microbial eukaryotes among different sampling conditions [11, 13–15]. The gene inventory conducted in all of our samples showed that 68% (64) of the 94 genes inventoried were present (S2 Table). Gene clustering of the 64 detected DEGs was performed using a transformed, normalized counts from variance stabilizing transformation generated in DESeq2 package (Fig 4). The genes were categorized based on the role they play in sexual development. These include meiosis, plasmogamy, karyogamy and other sexual and developmental related processes (Fig 4, S2 Table). Results of the DGE analysis for all these genes were shown in supplementary S2 Table with the log2 FC and adjusted p-value for each detected gene between single and fused conditions. We also included the results from comparisons that included an intermediate stage to explore our data in depth.

**Figure 4.**
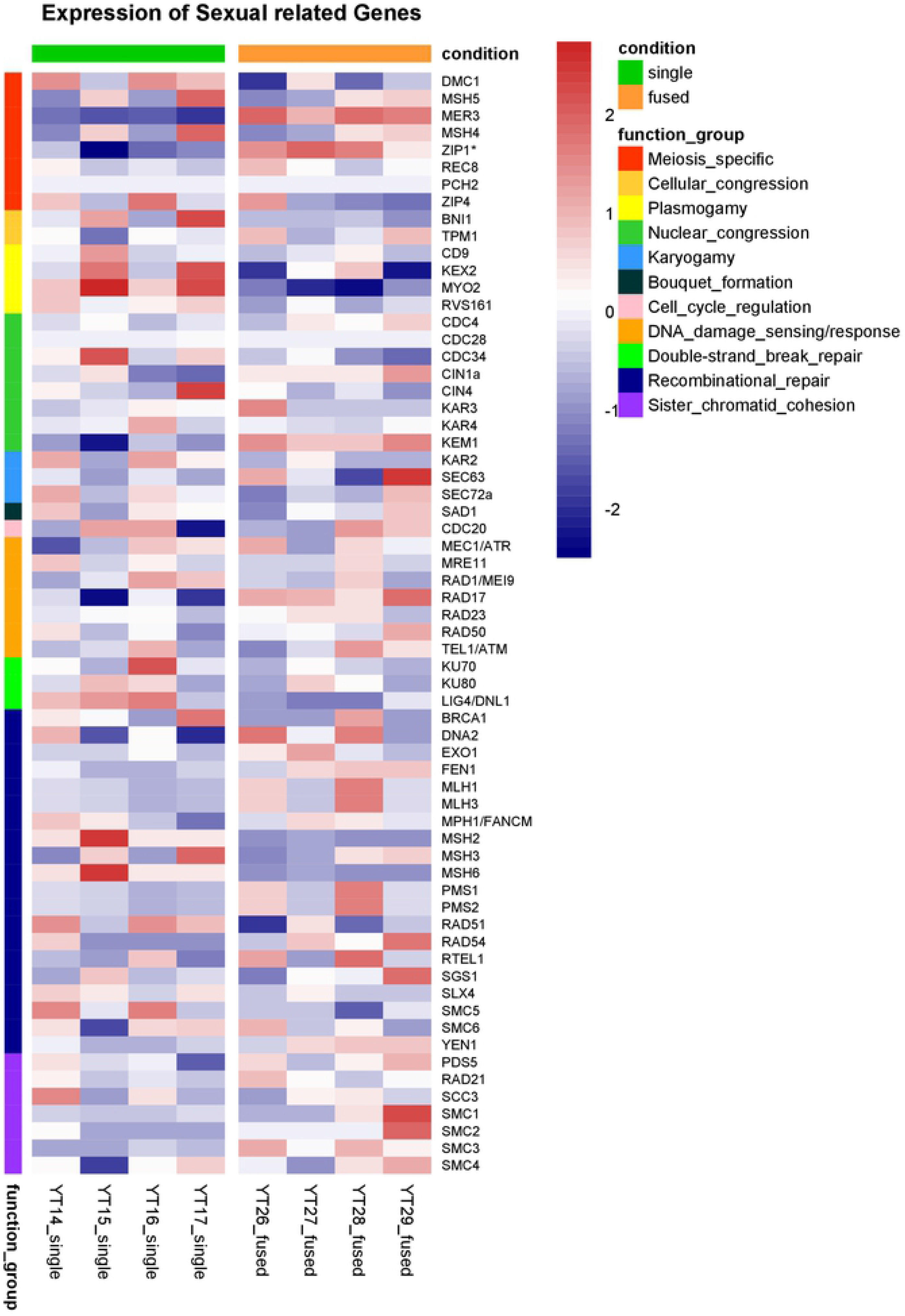
Heatmap of 64 genes (meiosis and sexual related) grouped according to their functional categories in single and fused samples. The heatmap was generated by centering the values across samples and thus shows the deviation of each gene in each sample using the data set from DESeq2 package. The color scale from red (highly expressed) to blue (low expression) represents the transformed, normalized counts from variance stabilizing transformation. Functional categories are shown in the color-coded annotation bar.

Among all the sexual related genes available for expression examination, 33 of the 64 genes analyzed were upregulated in fused cells, while single cells had 30 of these genes upregulated (see S2 Table for adjusted pvalue for each gene). One of the meiosis specific genes, Pch2, had no result due to its low expression (S2 Table). Among the 33 upregulated genes in fused samples, six were significant with threshold of adjusted pvalue of 0.05 (S2 Table). These include genes that are involved in meiosis (Mer3 and Zip1-like), Nuclear congression (Kem1), DNA damage sensing (Rad17) and Sister chromatid cohesion (Smc1/3) (Tables 1, S2, Fig 4). In single samples, five of the 30 upregulated genes were significant (S2 Table), which includes cellular congression (Bni1), plasmogamy (Myo2), double-strand break repair (Lig4/Dnl1) and recombinational repair (Smc5, Msh2, Msh6) (Fig 4, S2 Table). When looking into the expression for each functional category, all four genes involved in plasmogamy and some genes involved in double-strand break repair had upregulation in single condition samples, albeit most of them didn’t pass the significance threshold (Fig 4, S2 Table). Upregulation of genes in recombinational repair and sister chromatid cohesion categories were more prominent in fused samples, most of them with padj >0.05 (S2 Table). Expression pattern of some detected meiosis genes (e.g. Pch2) and other genes in S2 Table could not be analyzed due to low expression among samples or its lack in the reference transcriptome (e.g. Spo11). In general, we saw an overall trend that differentially expressed (upregulated) genes in the fused condition play an important role in sexual development. However, some meiosis specific genes such as Msh4/5 and Dmc1 were observed to be upregulated, without statistical significance, in some of the single cell samples (S2 Table).

**Table 1.**
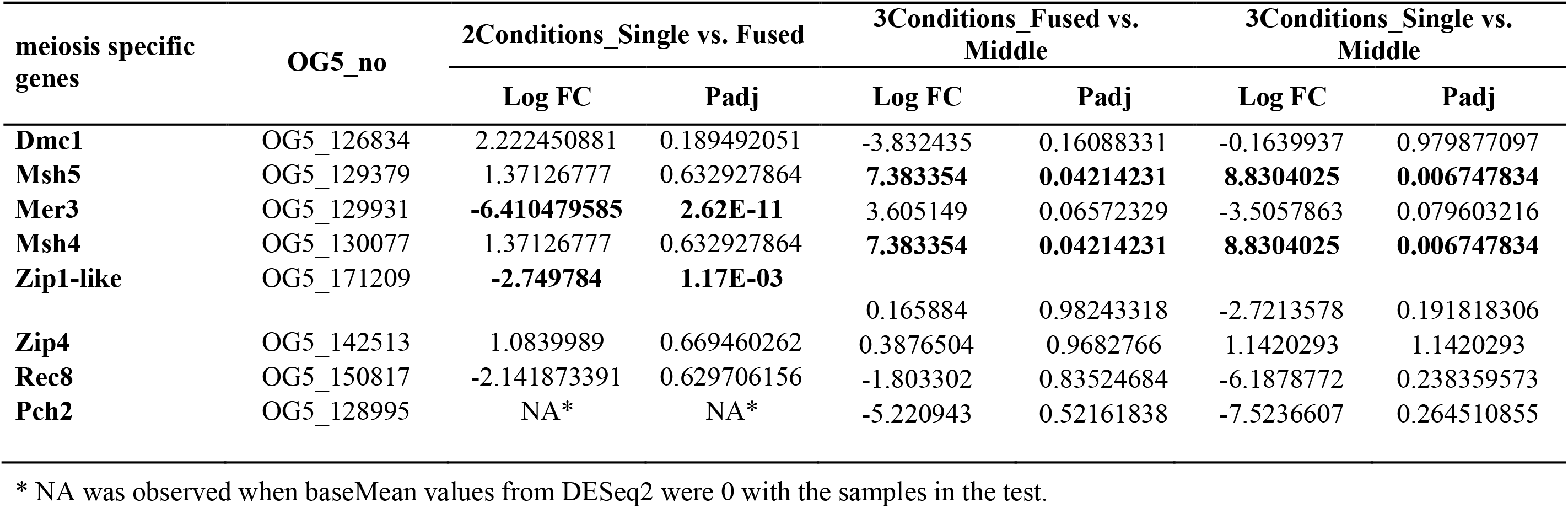
Expression results of meiosis specific genes in different group comparisons in terms of Log FC (bold significant) and Padj values.

### Functional annotation of DEGs

Blast2GO was applied to perform functional annotation of the two sets of DEGs identified among the single (521 genes) and fused (360 genes) samples. Three GO categories (Molecular function, Biological process, Cellular components) were inferred by mapping the sequence information to existing annotation sources. Based on these analyses, we were able to identify key functional categories that can be used to inform about sexual development in *Cochliopodium* (S7-S9 Figs). GO category representations in fused samples were dominated by nucleic acid (DNA) metabolism, nucleus and associated nuclear components (chromosome, nuclear pore, nucleolus), RNA processing, ATP binding and helicase activities. These are based on the most frequent GO terms and sequence distribution for each GO category (S7-S9 Figs). In addition, the fused samples consisted of some notable functional categories directly or indirectly related to sexual development (cell division) including ligase, histone acetylation, peroxisome fission (PHD-finger protein) and MCM complex (S7-S9 Figs). Enzyme Class (EC) distribution comparison of the two sets of DEGs showed that all the enzyme classes had more sequences in single samples than in fused samples except for the EC class Ligases (Fig 5). Various ligase enzymes were identified in fused samples including E3 ubiquitin-protein ligase previously shown to play a role in sexual development [47]. GO category representations in single samples were mostly dominated with metabolic activities (carbohydrate, lipid and protein), oxidation-reduction process, vesicle transport, cytoskeleton and cellular and intracellular movements (S7-S9 Figs). Among the top functional category representations in single samples are proteolysis from BP category and membrane from CC category (S7 Fig). Enrichment analysis showed that fused cells were abundant with GO categories of ATP binding and nucleus, while single cells were abundant with peptidase activity (S9 Fig).

**Figure 5.**
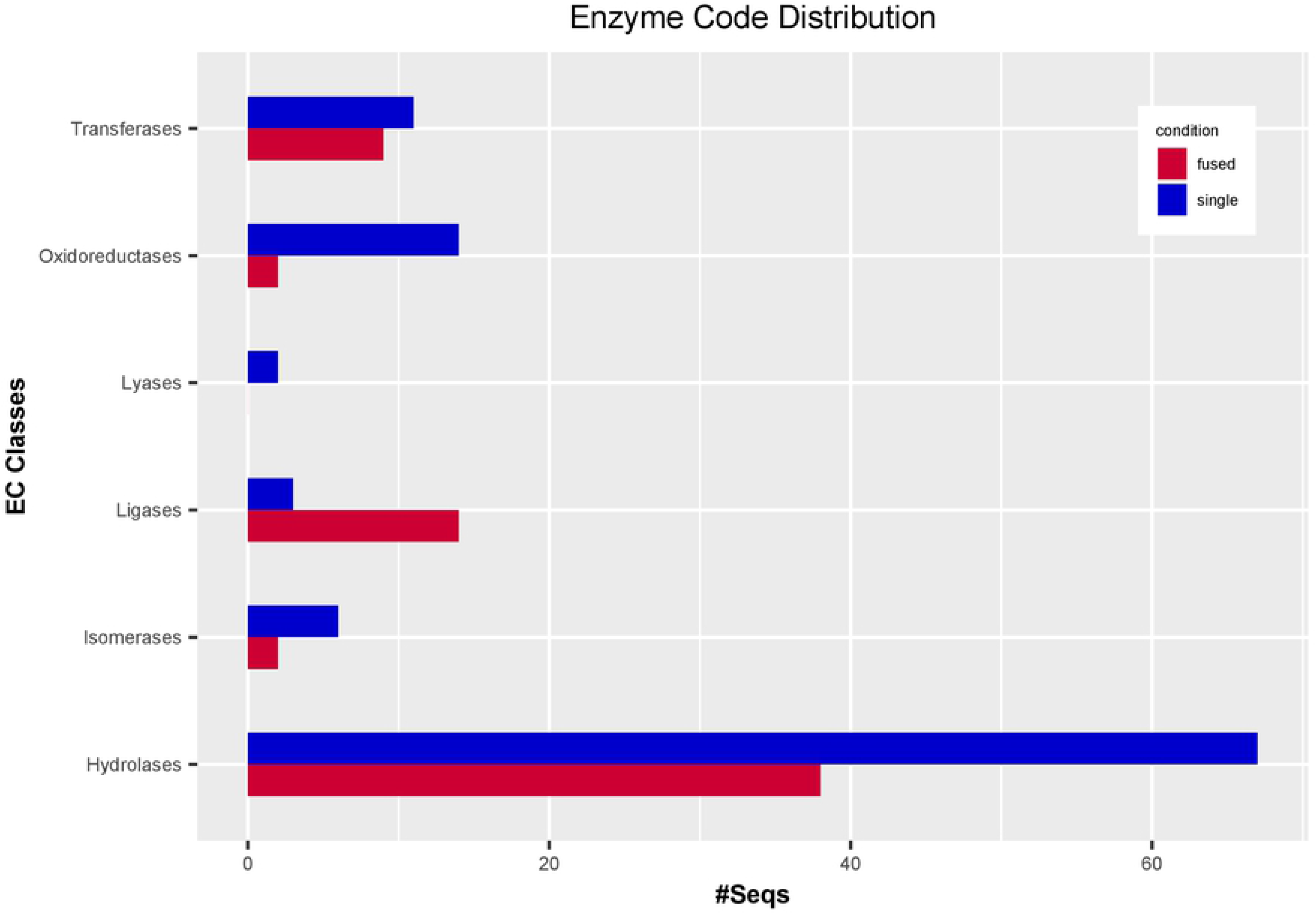
Enzyme Code (EC) distribution associated to the DEGs for single and fused group in terms of sequence numbers for six classes of EC derived from Blast2GO.

### Cytological detection of selected sexual DEGs

We performed RNA in situ hybridization (RNA-ISH) assay of selected genes representative of sexual development in the life cycle of *Cochliopodium*. RNA-ISH analysis gives both qualitative expression levels and spatial distribution of DEGs within intact cells. Our RNA-ISH results are mostly congruent to the DGE analysis of the selected genes. The meiosis genes, Mer3, Pch2 and Dmc1, and karyogamy gene (Kem1) were expressed mostly, but not exclusively, in fused cells (Figs 6-8A,B). The expression of Mer3 was the most prominent in fused cells at different stages pre-karyogamy (Fig 6A), during karyogamy (Fig 6B) and in polyploidy nucleus, post-karyogamy (Fig 6C). Mer3 was also expressed in single cells but at a much lower quantity (S10 Fig). The detection of Pch2 and Dmc1 was not as prominent as Mer3 (Fig 7). Pch2 was mostly detected in fused cells with multiple and polyploid nuclei (Fig 7C,D). Similarly, Dmc1 was detected mostly in fused samples but some single condition samples also expressed it (Fig 7A,B). The expression of Kem1 was noticeably visible around the nucleus periphery (Fig 8A,B), while the detection of Bni1 was not that prominent; it was expressed both in fused and single cells, but mostly in unfused/single cells (Fig 8C,D). We also attempted a colocalization (co-expression) experiment, but this experiment had many technical challenges. We were able to show that Mer3 and Dmc1 co-expressed in fused cells (S11 Fig).

**Figure 6.**
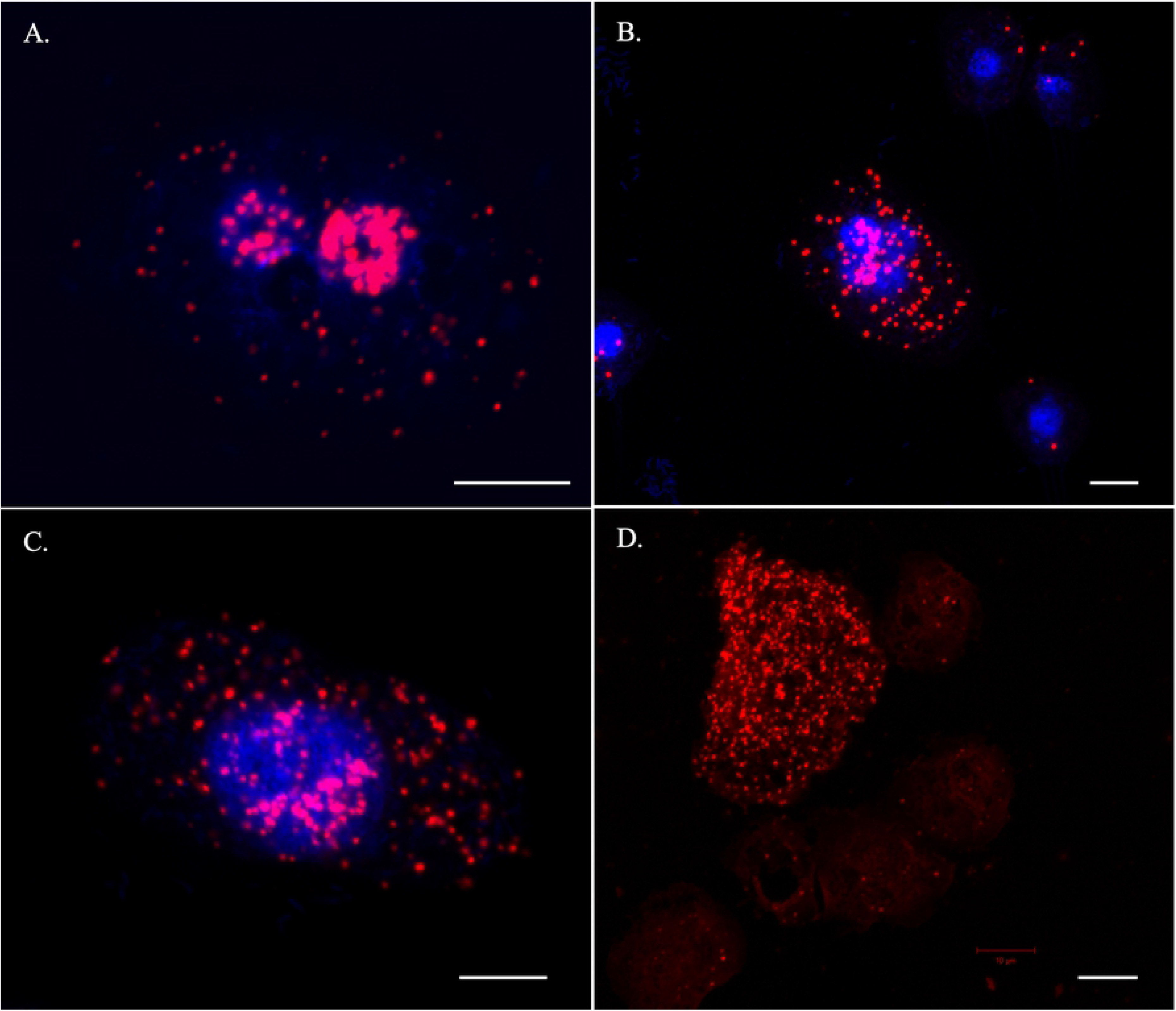
RNA-ISH staining of a meiosis specific gene, Mer3. The gene is expressed throughout the fused phases including pre-karyogamy (A), during-karyogamy (B) and post-karyogamy (C). D. Expression levels of Mer3 in a fused (top left) and non-fused amoeba cells (faint red staining). Red (Mer3) and Blue (DNA). Scale bar 10 μm.

**Figure 7.**
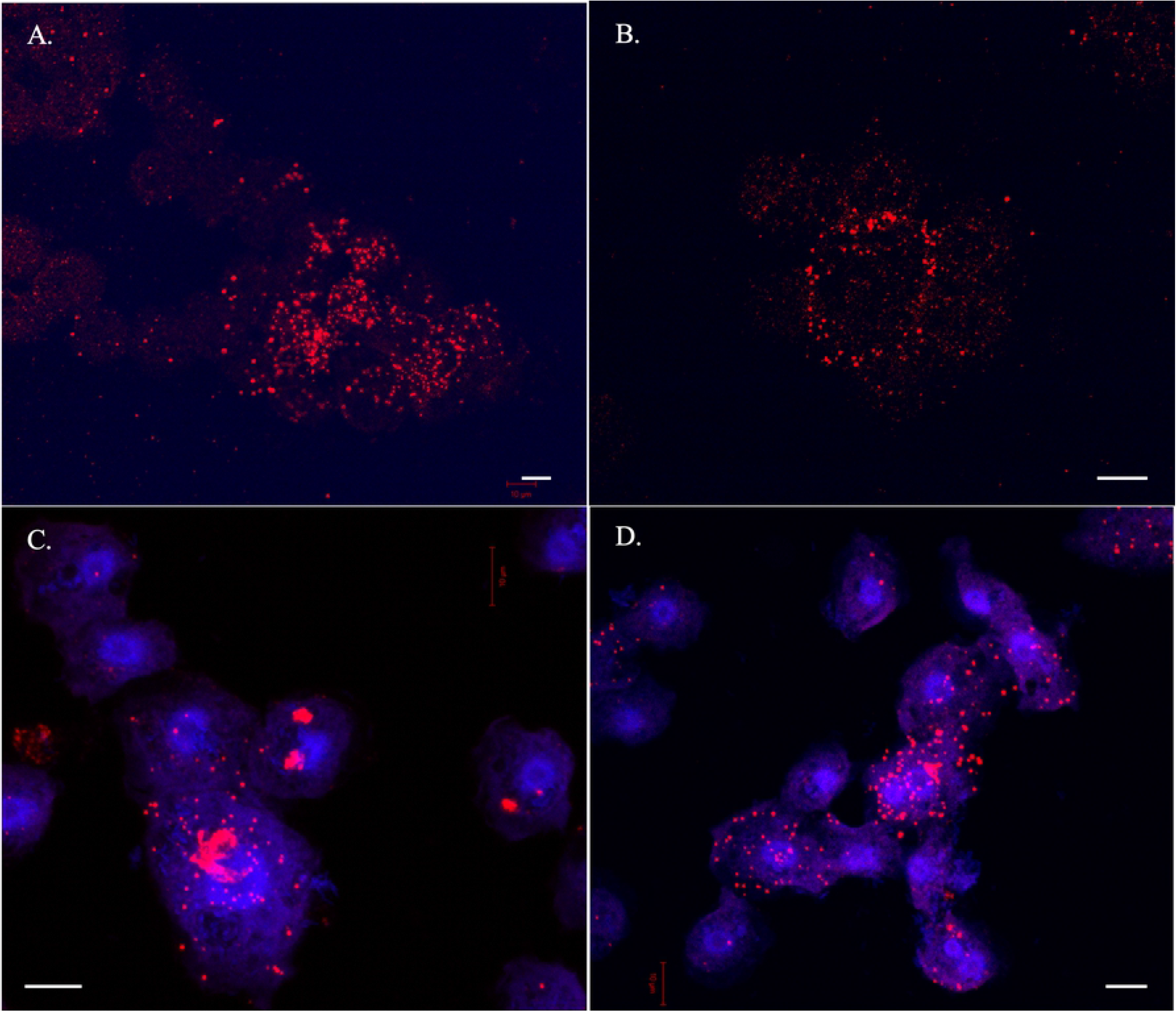
RNA-ISH staining of meiosis specific genes, Dmc1 (A-B) and Pch2 (C-D). Note high levels of expression of these genes in fused cells (arrows). Red (Mer3) and Blue (DNA). Scale bar 10 μm.

**Figure 8.**
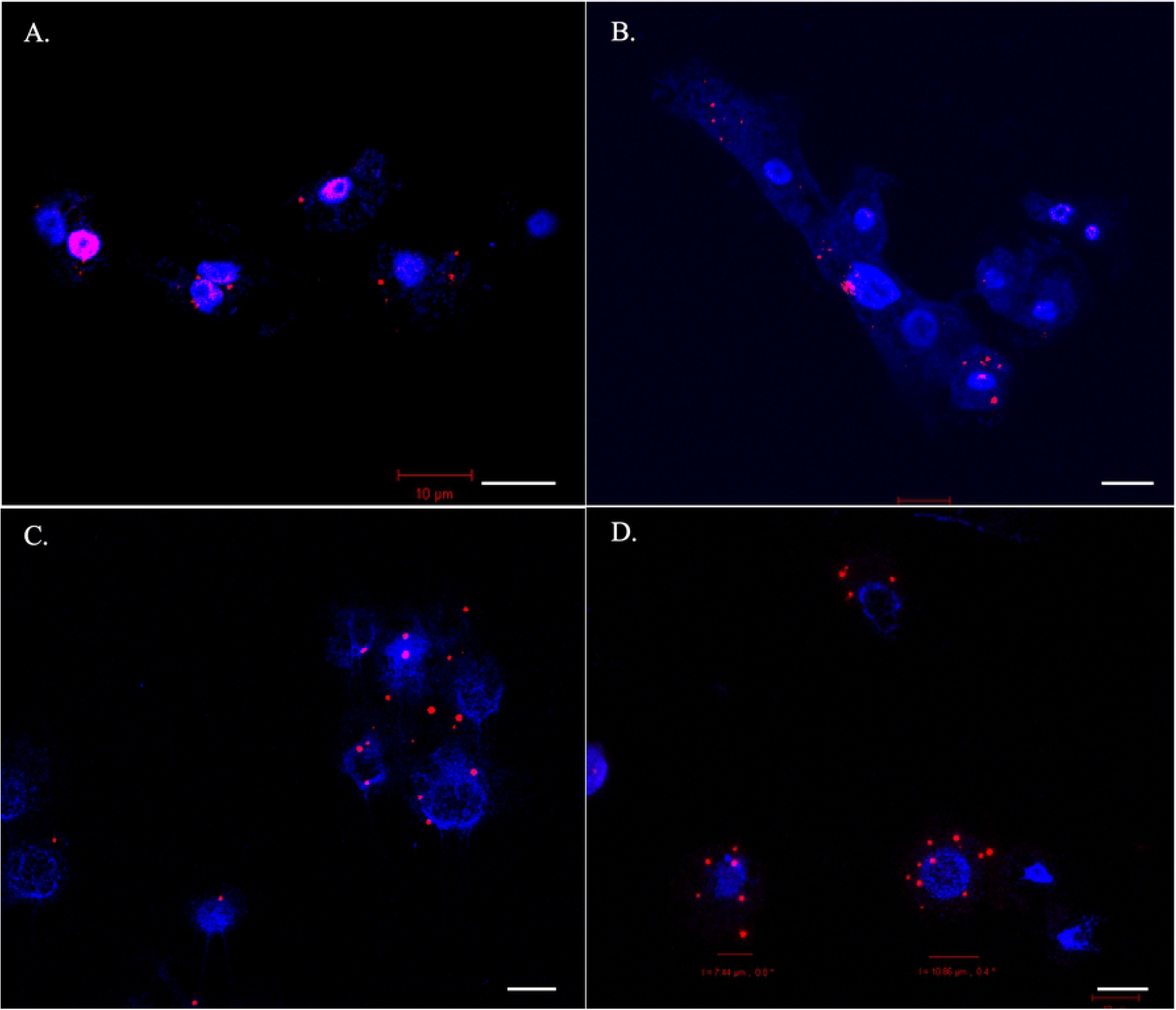
RNA-ISH staining of Karyogamy, Kem1 (A-B) and Plasmogamy Bni1 (C-D) genes. Note the localization of the karyogamy gene, Kem1, around nuclear periphery (A, arrows). Red (Mer3) and Blue (DNA). Scale bar 10 μm.

## Discussion

### Fused cells are sexual stage in *Cochliopodium*

The present study conclusively demonstrates that the fused cells in *Cochliopodium* both express and upregulate most of the meiosis and sexual related genes. Corroborative evidence from the cytological (RNA-ISH) study also provides spatial and temporal expression for some of these genes during the life cycle of *Cochliopodium* (Figs 6-8). Evidence for sex in a majority of amoebozoans was based on gene inventory of genomic or transcriptomic data, without supporting physical or stage specific DGE analysis [11]. Limited ultrastructural evidence for sex is available for a few non-model amoebae [22, 23, 28]. Most of the detailed genetic, DGE and cytological studies are limited to the model organism, *Dictyostelium discoideum* (e.g. [48–50]) and the pathogenic lineage, *Entamoeba* (e.g. [17, 20, 51–53]). However, both of these amoebae have dramatically different types of life cycle and could not be representative of the diverse life cycle observed in the supergroup: Amoebozoa. Our study is the first to report comprehensive DGE and cytological studies of sexual development in amoebae characterized by multiple cellular and nuclear fusions. The findings of this study have important implications in understanding molecular mechanisms of cryptic sexual processes in microbial eukaryotes that share similar behavior. Study of sexual processes in microbial eukaryotes has been curtailed mainly due to lack of observational evidence. This study demonstrates that *Cochliopodium* can serve as a model to understand some aspect of sexual processes shared among microbial eukaryotes due to its well-defined life cycle, tractability and suitability for genetic manipulation studies [16].

### Importance of cytological data in understanding DGE analysis in *Cochliopodium*

Our DGE analysis is largely corroborated by our RNA-ISH data for the selected sex genes. Even though most of our results from DGE and RNA-ISH were consistent with the expected sexual development of the amoeba, we observed some variations (albeit non-significant) in the DGE analysis among our samples (S2 Table). Among the variations noteworthy of mention include the detection and upregulation of meiosis specific genes in some single samples. These results prompted us to explore if sexual development in *Cochliopodium* occurs progressively with some intermediate stage. Our RNA-ISH results revealed some important clues that can help understand the observed variations. As shown in our three-sample condition analysis (S2-6 Figs) sexual development likely occurs in progressive fashion in *Cochliopodium* with some single cells expressing sex genes. These single cells are likely preparing to enter the sexual stage and can be considered intermediates. This observation is supported by the detection of meiosis specific genes in our RNA-ISH staining in some unfused uninucleate cells, although these genes were consistently detected in higher numbers in fused samples (Fig 6B, S10 Fig). All of our data sources are based on transcriptome (RNAseq), the detection of these genes in unfused cells, possibly entering sexual stage, might also be attributed to the time lag between transcription and translation.

The distinction between single and fused cells would have been more apparent if we used proteomic data. Immunostaining of meiosis protein could give substantial informative results pertaining to the actual stage of the amoeba. Our attempt to immunostain proteins encoding SPO11 and DMC1 using commercially available antibodies was not successful. Sex genes are among the fastest evolving genes in eukaryotes [11, 14, 54]. Some sex genes that serve similar function have evolved far beyond recognition making conventional sequence homology comparison impractical. These genes can only be identified through structural domain comparison or their localization in cytological studies (see [54]). Sex genes show great divergence even among closely related species. For example, the RNA-ISH probes designed for *Cochliopodium* did not yield good results in closely related species of amoebae (data not show). Proteomic work of sex genes is critical for determining sexual stages by providing physical (structural) localization of sexual stages as it occurs (e.g., synaptosomal complexes). However, this will require development of species-specific antibodies, which is challenging due to lack of genome data and the associated cost and time required to develop species-specific antibodies. With genome sequencing of many amoebae underway, a more comprehensive proteomic and gene manipulation experiments will be feasible in the near future, which will play an integral role in unraveling the various molecular mechanism of sexual strategies employed in the supergroup.

Another variation in DGE analysis relates to upregulation of some meiosis genes in single samples (Tables 1, S1, Fig 4). Although the upregulations of these genes (Msh5, Msh4 and Dmc1) were not significant (S2 Table), their detection in single samples requires an explanation. As described above our single samples data come from 5-10 cells and hence it is likely that our single samples might have included a mix of cells among which some cells were preparing to enter sexual stage. Our RNA-ISH data showed that single cells to some extent were observed to express meiosis genes (Figs 6B, S10). These single cells cannot be distinguished by cell size alone in live cultures and might partly explain the discrepancies between our DGE and RNA-ISH results. The detection of lowly expressed genes (e.g., Pch2) and a meiosis gene (Dmc1), shown to be upregulated in DGE analysis in single cells, were seen in our RNA-ISH to be more abundant in fused cells compared to in single cells (Figs 6-7, S10). Our RNA-ISH results are consistent with the expected life cycle of the amoeba. Another possible explanation for the observed discrepancies between DGE and RNA-ISH could be explained by the high adjusted pvalue (18.9 %) of Dmc1, which can be interpreted as false positive upregulation [44]. The cytological data greatly helped us to interpret DGE results and overcome some of the practical limitations of cell stage identification in *Cochliopodium*. Below we highlight the role of keys genes, differentially expressed (detected) in our samples, in the sexual development of *Cochliopodium*.

### DGE and cytological evidences point to the main crossover pathway employed in *Cochliopodium*

One of the most significantly upregulated DEGs in fused samples is Mer3 (Table 1, Fig 4). This gene is a meiosis specific gene and a member of the ZMM group described in budding yeast [55, 56]. Mer3 together with other members ZMM genes (Zip1-4, Msh4 and Msh5) play a crucial role in sexual organisms by providing a link between recombination and synaptonemal complex (SC) assembly during meiosis [57, 58]. Mer3 is among the highly expressed genes and was easily detected in cytological RNA-ISH staining in higher quantities (Fig 6). It was expressed at various phases of the fused cells including prior, during and post-nuclear fusion (karyogamy, Fig 6), which is indicative of its critical role in sexual development of *Cochliopodium*. Mer3, a highly evolutionary conserved DNA helicase, is involved in ZMM genes dependent interference cross-over (class I cross-over (CO) pathway), where double-Holliday junctions are preferentially resolved toward CO. Interference in this pathway prevents nearby crossovers from occurring thereby ensuring widely spaced crossovers along chromosomes [59, 60]. Most of the genes reported to play role in the class I CO pathway were either differentially expressed or detected in fused cell samples. These include meiosis specific ZMM genes (Zip4, Msh4 and Msh5) and other genes not specific to meiosis, including Mlh1-3, Exo1 and Sgs1 (Fig 4, S1 Table). We also found a significant DEG, Zip1-like, gene, which is one of the most important ZMM group members that forms the central component of SC [61]. However, the reference sequences of Zip1 gene retrieved from OrthoMCL database were few and quite divergent, which created a challenge for accurate homology assessment of this gene in our samples. Our homology search result for this gene in HMMER passed the inclusion threshold (HMMER c-evalue e-6) with a matching length of 414bp but failed the Blast (evalue e-7) threshold. HMMER is designed to detect remote homologs and takes into account shared protein domains in its search algorithm [45]. Based on this result, we decided to report the expression of Zip1-like genes from our transcriptome, which showed significant upregulation in fused cells (Table 1). However, this requires further investigation by searching similar genes in related species and acquisition of the full length of the gene in the genome of *Cochliopodium,* currently unavailable. Most of the ZMM group genes were described from budding yeast, similar genes playing the same role are known in plants and animals [62, 63]. It should be noted that the Zip1 gene of the budding yeast and other eukaryotes shares little sequence homology; however, their proteins share biochemical/structural similarities and localize in SC [54]. Given these observed divergences and the significant upregulation of the Zip1-like gene in *Cochliopodium* (Table 1), further analysis is needed to determine the identity and role of this gene in sexual development of the amoeba.

The expression and detection of Mer3 and most of ZMM group genes suggests that *Cochliopodium* predominantly employs class I CO and synaptonemal complexes for meiotic recombination. We also detected Pch2, a protein reported to play a regulatory role associated with other meiosis specific genes such as Zip1 and Hop1 (not detected here) in the maintenance of synaptonemal complex organization [64, 65]. Pch2 is one of the lowly expressed genes in our DGE analysis but was shown in our RNA-ISH data to localize more in fused samples (Fig 7C,D). Class I CO is a common pathway eukaryotes [54, 58], however, the exact mechanism in which class I CO pathway and SC operate in an amoeba characterized by multiparental nuclear fusion awaits further investigation. We also detected Mus81, one of the genes involve in non-interference class II CO pathway (S2 Table). Even though Mus81 is rarely detected in *Cochliopodium* samples, its presence indicates that class II CO might be used as alternative crossover pathway in this amoeba.

### Significance of cohesin complex in multiparental nuclear fusion

Most of the genes comprising the cohesin complex genes, Smc1-4 and Rad21 as well as the meiotic specific cohesin subunit, Rec8, that are essential in sister chromatid cohesion and their faithful separation, were expressed in fused cells [66–68]. Understanding the functionality and molecular mechanism of cohesin proteins is particularly of interest for further investigation in *Cochliopodium*. Cellular fusion in *Cochliopodium* is followed by mutiparental nuclear fusion and likely involves mixing of chromosomes from multiple individuals [16]. In conventional mitosis and meiosis, cohesins hold sister chromatids together before anaphase during cell division to ensure equal distribution of chromatids to dividing daughter/gamete cells. Any failure in cohesins results in aneuploidy, which has drastic health and viability consequences [69]. *Cochliopodium* presents an atypical parasexual process where fusion can result in over 30 nuclei in one large fused plasmodium. Investigation of how *Cochliopodium* sorts such a large number of multiparental chromosomes precociously during subsequent fission of fused cells will likely unravel a highly evolved or novel mechanism for the roles of cohesins.

### Other meiosis and sex related genes

We detected several meiosis and sex-related genes that play a role in mismatch repair, initiation of double-strand break (DSB) and their repair as well as homologous recombination (Fig 4, S2 Table). Although these genes are detected in our DGE analysis, their patterns of expression were neither significant nor consistent in our samples (S2 Table). For example, genes involved in Homologous recombination (HR), Rad51 and Dmc1 (meiosis specific), were expressed and upregulated in our single samples. Although Dmc1 was shown to localize with Mer3 in our fused samples (S14 Fig), the temporal expression of these genes needs further investigation. Both of these genes code for recombinases that play critical role in DNA lesions repair including double-strand breaks (DSBs), single-strand DNA gaps and interstrand crosslinks during meiosis [70]. Interestingly, genes involved in the alternative DSBs repair mechanism, nonhomologous DNA end joining (NHEJ), were all upregulated in single samples. Proteins involved in NHEJ pathway including KU70 and KU80 are operational during the vegetative stage (G0/G1 phases of the cell cycle) when sister chromatids are not available [71]. These genes are known to be downregulated during meiosis [72], which is consistent with the DGE results of our fused samples (Fig 4, S2 Table). Lastly, one of the key meiosis genes, Spo11, that initiates meiosis by programed DSB was not detected in our DGE analysis. Spo11 is detected in several species of amoebae, but some amoebae (e.g., *Dictyostelium*) lack it in their genome [11, 12, 73]. Spo11 is likely one of the lowly expressed genes since we only detected it one time in one sample; and our attempt to detect it using RNA-ISH with a probe designed from a closely related species rendered no results.

### Other DEGs and Gene ontology (GO) provide information on the developmental and sexual stage of *Cochliopodium*

Additional DEGs and GO categories mirroring cellular physiology of the amoeba shed light on the life cycle of *Cochliopodium*. Genes engaged in plasmogamy and karyogamy were upregulated in single and fused samples, respectively (Fig 4, S2 Table). This result is consistent with the observed life cycle of *Cochliopodium,* where single cells entering a sexual stage would be expected to produce proteins enabling them to fuse, and expression of karyogamy genes in fused cells supports the observed nuclear fusion [16]. Among the most dominant DEGs in the GO analyses in single samples were cellular processes related to metabolic activity and vesicles mediated transportation (S7-S9 Figs). These are indicative of an active vegetative phase of our single samples. Particularly, detection of a large number of proteolytic enzymes indicates that the cells might be engaging in high protein metabolism (S7 Fig). The most dominate GO terms in fused samples included DNA metabolism, nucleus, ATP binding, PHD-Finger protein and ligases (S7-S9 Figs). These results can be interpreted as fused cells primarily engaged in ATP-mediated, DNA-related processes, likely reflecting their sexual nature. Among these, E3 ubiquitin ligase [47] and PHD-Finger [74] have been implicated to play a role in sexual development of plants. E3 ubiquitin ligase has a similar functional domain (i.e., RING-finger to Zip3), a meiosis specific gene and a member of the ZMM group known to play roles in crossover and SC assembly [54, 75]. While this finding will require in depth analysis of these genes, the overall physiological and cellular processes of the GO analysis lend support to the sexual nature of the fused samples.

### Understanding sexuality in Amoebozoa and future directions

We are just starting to scratch the surface of the enormous life cycle diversity in the supergroup Amoebozoa. The current study and previous studies indicated that members of Amoebozoa might use various mechanisms of sexual pathways that reflect their life cycle and behavior [11, 76]. For example, *Dicytostelium* has lost Spo11 but most amoebae seem to possess this gene [11, 12, 73]; this observation indicates the existence of variation in the mechanism of meiosis initiation in amoebozoans. A closer look of the known (yet to be described) diversity of the amoebozoans will likely uncover even additional, and novel, forms of life cycle and sexual strategies in this supergroup. Another layer of complexity in understanding sexuality in amoebae is the observation that some amoebae that display no signs of sexual activity constitutively, nonetheless express meiosis specific genes in actively growing cultures. Such an example is *Acanthamoeba*, a well-studied amoebozoan lineage that show no evidence of fusion or other sexual like behavior during its life cycle [76]. *Acanthamoeba* and other amoebae are reported to change their ploidy during their life cycle [77–80]. This observation has led to a suggestion that amoebae might still be asexual lineages but have evolved to use meiosis genes to perform recombination through other means such as polyploidization and gene conversion [81]. Meiosis genes have been used as a blueprint for sex, however, it has been suggested that detection of meiosis genes should not be interpreted as evidence for sexual reproduction [82]. More investigation is required to unravel the nature and roles of meiosis genes in the life cycle of amoebzoans.

Despite the progress we have made in the current study, many fundamental questions about the sexual nature and details of the life cycle of *Cochliopodium* remain unknown. Mating types, the nature of multiparental inheritance and polyploidization in *Cochliopodium* are still poorly elucidated. *Cochliopodium* has been shown to fuse even in monoclonal cultures, though fusion frequency was lower compared to mixed cultures (S12 Fig). Mating types are known in *Dictyostelium* and other related amoebae [49, 83–85], further investigation is required to examine if fusion in *Cochliopodium* occurs randomly or among compatible mating types. The mechanism of inheritance involving multiple partners is a challenge to the classic mechanism of inheritance in dimorphic systems – well known in various eukaryotes. Triparental inheritance involving more than two mating types has been reported in *Dictyostelium* [50]. Further investigation using genome data, gene manipulation and cell biology studies are required to elucidate these fundamental questions in *Cochliopodium* and other amoebae showing similar life cycle behaviors.

## Acknowledgments

We would like to thank James T. Melton III, Fiona Wood, Stephen Kioko and Tori Concepcion for technical assistance during data collection. O. Roger Anderson is thanked for his useful comments and edits on the manuscript.

## Supporting information

**Figure S1**. PCA plot of the 18 samples in the two-experiment condition (single vs. fused). PCA data for the plot were the transformed normalized counts of each sample generated from DESeq2.

**Figure S2.** PCA plot of a three-condition experiment (single, fused and intermediate) each represented by 3 replicates.

**Figure S3.** Venn diagram of the number of DEGs (Padj < 0.05) within the three-condition experiment, showing shared DEGs between experimental conditions: single vs. intermediate, single vs. fused, and intermediate vs. fused.

**Figure S4.** Clustered heatmap of all the 738 DEGs identified between each of the 2-stage comparisons in the three-condition experiment with a total 9 replicates. The color scale from red (highly expressed) to blue (low expression) represents the transformed, normalized counts from variance stabilizing transformation.

**Figure S5.** Heatmap of the 64 genes (meiosis and sexually related) grouped according to their function categories for the three-condition experiment. The heatmap was generated by centering the values across samples and thus shows the deviation of each gene in each sample using the data set from DESeq2 package. The color scale from red (highly expressed) to blue (low expression) represents the transformed, normalized counts from variance stabilizing transformation. Function categories were shown in the annotation bar.

**Figure S6.** Heatmap of the meiosis specific genes grouped according to their function categories for the three-condition experiment: single, fused, intermediate.

**Figure S7.** Direct Go Count plots showing the most frequent GO terms from each set of DEGs identified from the two-condition experiment for each GO category (Biological process (BP), Cellular components (CC), and Molecular function (MF)). The plots were generated from Blast2GO without taking account of the GO hierarchy. The left panel shows the results for 360 DEGs upregulated in fused samples and the right panel shows the results for 521 DEGs upregulated in single samples.

**Figure S8**. Multilevel pie chart showing the sequence distribution by GO categories. These were generated with the lowest node per branch of the combined GO graph from Blast2GO for each of the three GO categories (Molecular function, Biological process, Cellular components) with sequence number and percentage. The left panel shows the results for 360 DEGs upregulated in fused samples and the right panel shows the results for 521 DEGs upregulated in single samples.

**Figure S9**. Enrichment analysis of the 2 sets of DEGs from Blast2GO. (A) The test gene set was for the 360 DEGs upregulated in fused samples. (B) The test gene set was for the 521 DEGs upregulated in single samples. In both tests, all 881 DEGs were set as the reference set.

**Figure S10**. RNA-ISH staining of a meiosis specific gene, Mer3, showing the distribution of Mer3 expression in single, fused and possibly intermediate cells (A-C). Note that the variation in expression levels among single uninucleate cells. In (A) single cells seem to be expressing less Mer3 than single cells in (B). Mer3 is consistently expressed in higher quantity in fused cells (A and B, arrows). Red (Mer3) and Blue (DNA). Scale bar 10 μm.

**Figure S11.** RNA-ISH staining showing the co-expression of two meiosis specific genes, Mer3 (red) and Dmc1 (green) in fused cells.

**Figure S12.** Fusion rate measured by average size between monoclonal (broken) and mixed (solid) cultures.

## Reference

1. Lahr D.J., Parfrey L.W., Mitchell E.A., Katz L.A., Lara E. 2011 The chastity of amoebae: re-evaluating evidence for sex in amoeboid organisms. Proceedings Biological sciences / The Royal Society 278(1715), 2081–2090. (doi:10.1098/rspb.2011.0289).

2. Wenrich D.H. 1954 Sex in Protozoa: a comparative review. Washington, DC: AAAS.

3. Williams G.C. 1975 Sex and evolution. Princeton, Princeton University Press.

4. Cavalier-Smith T. 2002 Origins of the machinery of recombination and sex. Heredity 88, 125–141.

5. Margulis L., Sagan D. 1985 Evolutionary origins of Sex. In Oxford Surveys in Evolutionary Biology (eds. Dawkins R., Ridley M.), pp. 30–47. Oxford, England, Oxford University Press.

6. Smith J.M. 1986 Evolution: contemplating life without sex. Nature 324(6095), 300–301. (doi:10.1038/324300a0).

7. Smith J.M., Szathmáry E. 1995 The Major Transitions in Evolution., W. H. Freeman and Company; 149 p.

8. Raikov I.B. 1995 Meiosis in protists: recent advances and persisting problems.. Eur J Protistol 31, 1–7.

9. Fallis A.M. 1965 Protozoan Life Cycles. Am Zool 5, 85–94.

10. Bell G., Koufopanou V., Harvey P.H., Partridge L., Southwood R. 1997 The architecture of the life cycle in small organisms. Phil Trans R Soc Lond B, 33281–33289. (doi:http://doi.org/10.1098/rstb.1991.0035).

11. Tekle Y.I., Wood F.C., Katz L.A., Ceron-Romero M.A., Gorfu L.A. 2017 Amoebozoans Are Secretly but Ancestrally Sexual: Evidence for Sex Genes and Potential Novel Crossover Pathways in Diverse Groups of Amoebae. Genome Biol Evol 9(2), 375–387. (doi:10.1093/gbe/evx002).

12. Hofstatter P.G., Brown M.W., Lahr D.J.G. 2018 Comparative Genomics Supports Sex and Meiosis in Diverse Amoebozoa. Genome Biol Evol 10(11), 3118–3128. (doi:10.1093/gbe/evy241).

13. Wood F.C., Heidari A., Tekle Y.I. 2017 Genetic Evidence for Sexuality in Cochliopodium (Amoebozoa). J Hered 108(7), 769–779. (doi:10.1093/jhered/esx078).

14. Malik S.B., Pightling A.W., Stefaniak L.M., Schurko A.M., Logsdon J.M., Jr. 2008 An expanded inventory of conserved meiotic genes provides evidence for sex in Trichomonas vaginalis. PloS one 3(8), e2879. (doi:10.1371/journal.pone.0002879).

15. Ramesh M.A., Malik S.-B., Logsdon J.M. 2005 A Phylogenomic Inventory of Meiotic Genes: Evidence for Sex in Giardia and an Early Eukaryotic Origin of Meiosis. 15(2), 185.

16. Tekle Y.I., Anderson O.R., Lecky A.F. 2014 Evidence of parasexual activity in “asexual amoebae” Cochliopodium spp. (Amoebozoa): extensive cellular and nuclear fusion. Protist 165(5), 676–687. (doi:10.1016/j.protis.2014.07.008).

17. Mukherjee C., Clark G., Lohia A. 2008 *Entamoeba* shows reversible variation in ploidy under different growth conditions and between life cycle phases. PLoS neglected tropical diseases 2(8), e281.

18. Chang M.T., Raper K.B. 1981 Mating types and macrocyst formation in Dictyostelium. Journal of bacteriology 147(3), 1049–1053.

19. Goodfellow L.P., Belcher J.H., Page F.C. 1974 A light- and electron-microscopical study of Sappinia diploidea, a sexual amoeba. Protistologica 2, 207–216.

20. Schensnovich V.B. 1967 [On the life cycle of Entamoeba histolytica]. Meditsinskaia parazitologiia i parazitarnye bolezni 36(6), 712–715.

21. Tekle Y.I., Anderson O.R., Lecky A.F., Kelly S.D. 2013 A New Freshwater Amoeba: Cochliopodium pentatrifurcatum n. sp. (Amoebozoa, Amorphea). The Journal of eukaryotic microbiology 60(4), 342–349. (doi:10.1111/jeu.12038).

22. Aldrich H.C. 1967 The ultrastructure of meiosis in three species of Physarum. Mycologia 59(1), 127–148.

23. Furtado J.S., Olive L.S. 1971 Ultrastructural evidence of meiosis in Ceratiomyxa fruticulosa.. Mycologia 63(2), 413–416.

24. Haskins E.F., Hinchee A.A., Cloney R.A. 1971 The occurrence of synaptonemal complexes in the slime mold Echinostelium minutum de Bary. J Cell Biol 51(3), 898–903. (doi:10.1083/jcb.51.3.898).

25. Erdos G.W., A.W. N., K.B. R. 1972 Fine structure of macrocysts in *Polysphondylium violaceum*. Cytobiologie 6, 351–366.

26. Erdos G.W., Raper K.B., Vogen L.K. 1975 Sexuality In Cellular Slime-Mold Dictyostelium-Giganteum. Proceedings Of The National Academy Of Sciences Of The United States Of America 72(3), 970–973.

27. Szabo S.P., O’Day D.H., Chagla A.H. 1982 Cell fusion, nuclear fusion, and zygote differentiation during sexual development of Dictyostelium discoideum. Dev Biol 90(2), 375–382. (doi:10.1016/0012-1606(82)90387-6).

28. Mignot J.-P., Raikov I.B. 1992 Evidence for meiosis in the testate amoeba *Arcella*. Journal of Eurkaryotic Microbiology 39(2), 287–289.

29. Martin G.W., Alexopoulos C.J. 1969 The Myxomycetes. Iowa City, University of Iowa Press; 201 p.

30. Schaudinn F. 1899 Untersuchungen über den Generationswechsel von Trichosphaerium sieboldi. Schn Abh Königl Preuss Akad Wiss, Berlin,, 93 p.

31. Anderson O.R., Tekle Y.I. 2013 A Description of Cochliopodium megatetrastylus n. sp. Isolated from a Freshwater Habitat. Acta Protozool 52, 55–64. (doi:doi:10.4467/16890027AP.13.006.1085).

32. Melton III J.T., M. S., F.C. W., Collins S.J., Y.I. T. 2019 Three New Freshwater Cochliopodium Species (Himatismenida, Amoebozoa) from the Southeastern United States. Journal of Eukaryotic Microbiology 0, 1–13. (doi:https://doi.org/10.1111/jeu.12764).

33. Davoli T., de Lange T. 2011 The causes and consequences of polyploidy in normal development and cancer. Annu Rev Cell Dev Biol 27, 585–610. (doi:10.1146/annurev-cellbio-092910-154234).

34. Duncan A.W., Hickey R.D., Paulk N.K., Culberson A.J., Olson S.B., Finegold M.J., Grompe M. 2009 Ploidy reductions in murine fusion-derived hepatocytes. PLoS Genet 5(2), e1000385. (doi:10.1371/journal.pgen.1000385).

35. Tekle Y.I., Wood F.C. 2018 A practical implementation of large transcriptomic data analysis to resolve cryptic species diversity problems in microbial eukaryotes. BMC evolutionary biology 18(1), 170. (doi:10.1186/s12862-018-1283-1).

36. Bankevich A., Nurk S., Antipov D., Gurevich A.A., Dvorkin M., Kulikov A.S., Lesin V.M., Nikolenko S.I., Pham S., Prjibelski A.D., et al. 2012 SPAdes: a new genome assembly algorithm and its applications to single-cell sequencing. J Comput Biol 19(5), 455–477. (doi:10.1089/cmb.2012.0021).

37. Bray N.L., Pimentel H., Melsted P., Pachter L. 2016 Near-optimal probabilistic RNA-seq quantification. Nat Biotechnol 34(5), 525–527. (doi:10.1038/nbt.3519).

38. Tekle Y.I., Anderson O.R., Katz L.A., Maurer-Alcala X.X., Romero M.A., Molestina R. 2016 Phylogenomics of ‘Discosea’: A new molecular phylogenetic perspective on Amoebozoa with flat body forms. Molecular phylogenetics and evolution 99, 144–154. (doi:10.1016/j.ympev.2016.03.029).

39. Tekle Y.I., Wood F.C. 2017 Longamoebia is not monophyletic: Phylogenomic and cytoskeleton analyses provide novel and well-resolved relationships of amoebozoan subclades. Molecular phylogenetics and evolution 114, 249–260. (doi:10.1016/j.ympev.2017.06.019).

40. Wood F.C., Heidari A., Tekle Y.I. 2017 Genetic Evidence for Sexuality in Cochliopodium. Journal of Heredity. (doi:https://doi.org/10.1093/jhered/esx078).

41. Love M.I., Huber W., Anders S. 2014 Moderated estimation of fold change and dispersion for RNA-seq data with DESeq2. Genome biology 15(12), 550. (doi:10.1186/s13059-014-0550-8).

42. Robinson M.D., McCarthy D.J., Smyth G.K. 2010 edgeR: a Bioconductor package for differential expression analysis of digital gene expression data. Bioinformatics 26(1), 139–140. (doi:10.1093/bioinformatics/btp616).

43. Schurch N.J., Schofield P., Gierlinski M., Cole C., Sherstnev A., Singh V., Wrobel N., Gharbi K., Simpson G.G., Owen-Hughes T., et al. 2016 How many biological replicates are needed in an RNA-seq experiment and which differential expression tool should you use? RNA 22(6), 839–851. (doi:10.1261/rna.053959.115).

44. Benjamini Y., Hochberg Y. 1995 Controlling the false discovery rate: a practical and powerful approach to multiple testing.. J Royal Stat Soc Ser B 57(1), 449–518.

45. Eddy S.R. 2001 HMMER: Profile hidden markov models for biological sequence analysis. (Available from: http://hmmer.janelia.org/.

46. Gotz S., Garcia-Gomez J.M., Terol J., Williams T.D., Nagaraj S.H., Nueda M.J., Robles M., Talon M., Dopazo J., Conesa A. 2008 High-throughput functional annotation and data mining with the Blast2GO suite. Nucleic Acids Res 36(10), 3420–3435. (doi:10.1093/nar/gkn176).

47. Zuhl L., Volkert C., Ibberson D., Schmidt A. 2019 Differential activity of F-box genes and E3 ligases distinguishes sexual versus apomictic germline specification in Boechera. J Exp Bot 70(20), 5643–5657. (doi:10.1093/jxb/erz323).

48. Urushihara H., Muramoto T. 2006 Genes involved in Dictyostelium discoideum sexual reproduction. Eur J Cell Biol 85(9-10), 961–968. (doi:10.1016/j.ejcb.2006.05.012).

49. Bloomfield G., Skelton J., Ivens A., Tanaka Y., Kay R.R. 2010 Sex determination in the social amoeba Dictyostelium discoideum. Science 330(6010), 1533–1536. (doi:10.1126/science.1197423).

50. Bloomfield G., Paschke P., Okamoto M., Stevens T.J., Urushihara H. 2019 Triparental inheritance in Dictyostelium. Proc Natl Acad Sci U S A 116(6), 2187–2192. (doi:10.1073/pnas.1814425116).

51. De Cadiz A.E., Jeelani G., Nakada-Tsukui K., Caler E., Nozaki T. 2013 Transcriptome analysis of encystation in Entamoeba invadens. PloS one 8(9), e74840. (doi:10.1371/journal.pone.0074840).

52. Singh N., Bhattacharya A., Bhattacharya S. 2013 Homologous recombination occurs in Entamoeba and is enhanced during growth stress and stage conversion. PloS one 8(9), e74465. (doi:10.1371/journal.pone.0074465).

53. Stanley S.L., Jr. 2005 The Entamoeba histolytica genome: something old, something new, something borrowed and sex too? Trends in parasitology 21(10), 451–453. (doi:S1471-4922(05)00213-8 [pii] 10.1016/j.pt.2005.08.006).

54. Pyatnitskaya A., Borde V., De Muyt A. 2019 Crossing and zipping: molecular duties of the ZMM proteins in meiosis. Chromosoma 128(3), 181–198. (doi:10.1007/s00412-019-00714-8).

55. Jessop L., Rockmill B., Roeder G.S., Lichten M. 2006 Meiotic chromosome synapsis-promoting proteins antagonize the anti-crossover activity of sgs1. PLoS Genet 2(9), e155. (doi:10.1371/journal.pgen.0020155).

56. Borner G.V., Kleckner N., Hunter N. 2004 Crossover/noncrossover differentiation, synaptonemal complex formation, and regulatory surveillance at the leptotene/zygotene transition of meiosis. Cell 117(1), 29–45. (doi:10.1016/s0092-8674(04)00292-2).

57. Nakagawa T., Ogawa H. 1999 The Saccharomyces cerevisiae MER3 gene, encoding a novel helicase-like protein, is required for crossover control in meiosis. EMBO J 18(20), 5714–5723. (doi:10.1093/emboj/18.20.5714).

58. Lynn A., Soucek R., Borner G.V. 2007 ZMM proteins during meiosis: crossover artists at work. Chromosome research: an international journal on the molecular, supramolecular and evolutionary aspects of chromosome biology 15(5), 591–605. (doi:10.1007/s10577-007-1150-1).

59. Nakagawa T., Kolodner R.D. 2002 The MER3 DNA helicase catalyzes the unwinding of holliday junctions. J Biol Chem 277(31), 28019–28024. (doi:10.1074/jbc.M204165200).

60. Zakharyevich K., Tang S., Ma Y., Hunter N. 2012 Delineation of joint molecule resolution pathways in meiosis identifies a crossover-specific resolvase. Cell 149(2), 334–347. (doi:10.1016/j.cell.2012.03.023).

61. Sym M., Engebrecht J.A., Roeder G.S. 1993 ZIP1 is a synaptonemal complex protein required for meiotic chromosome synapsis. Cell 72(3), 365–378. (doi:10.1016/0092-8674(93)90114-6).

62. Higgins J.D., Sanchez-Moran E., Armstrong S.J., Jones G.H., Franklin F.C. 2005 The Arabidopsis synaptonemal complex protein ZYP1 is required for chromosome synapsis and normal fidelity of crossing over. Genes Dev 19(20), 2488–2500. (doi:10.1101/gad.354705).

63. de Vries F.A., de Boer E., van den Bosch M., Baarends W.M., Ooms M., Yuan L., Liu J.G., van Zeeland A.A., Heyting C., Pastink A. 2005 Mouse Sycp1 functions in synaptonemal complex assembly, meiotic recombination, and XY body formation. Genes Dev 19(11), 1376–1389. (doi:10.1101/gad.329705).

64. Chen C., Jomaa A., Ortega J., Alani E.E. 2014 Pch2 is a hexameric ring ATPase that remodels the chromosome axis protein Hop1. Proc Natl Acad Sci U S A 111(1), E44–53. (doi:10.1073/pnas.1310755111).

65. Zanders S., Sonntag Brown M., Chen C., Alani E. 2011 Pch2 modulates chromatid partner choice during meiotic double-strand break repair in Saccharomyces cerevisiae. Genetics 188(3), 511–521. (doi:10.1534/genetics.111.129031).

66. Watanabe Y. 2004 Modifying sister chromatid cohesion for meiosis. J Cell Sci 117(Pt 18), 4017–4023. (doi:10.1242/jcs.01352).

67. Klein F., Mahr P., Galova M., Buonomo S.B., Michaelis C., Nairz K., Nasmyth K. 1999 A central role for cohesins in sister chromatid cohesion, formation of axial elements, and recombination during yeast meiosis. Cell 98(1), 91–103. (doi:10.1016/S0092-8674(00)80609-1).

68. Peters J.M., Tedeschi A., Schmitz J. 2008 The cohesin complex and its roles in chromosome biology. Genes Dev 22(22), 3089–3114. (doi:10.1101/gad.1724308).

69. Chiang T., Duncan F.E., Schindler K., Schultz R.M., Lampson M.A. 2010 Evidence that weakened centromere cohesion is a leading cause of age-related aneuploidy in oocytes. Current biology: CB 20(17), 1522–1528. (doi:10.1016/j.cub.2010.06.069).

70. Krejci L., Altmannova V., Spirek M., Zhao X. 2012 Homologous recombination and its regulation. Nucleic Acids Res 40(13), 5795–5818. (doi:10.1093/nar/gks270).

71. Ahmed E.A., Sfeir A., Takai H., Scherthan H. 2013 Ku70 and non-homologous end joining protect testicular cells from DNA damage. J Cell Sci 126(Pt 14), 3095–3104. (doi:10.1242/jcs.122788).

72. Goedecke W., Eijpe M., Offenberg H.H., van Aalderen M., Heyting C. 1999 Mre11 and Ku70 interact in somatic cells, but are differentially expressed in early meiosis. Nat Genet 23(2), 194–198. (doi:10.1038/13821).

73. Bloomfield G. 2018 Spo11-Independent Meiosis in Social Amoebae. Annu Rev Microbiol 72, 293–307. (doi:10.1146/annurev-micro-090817-062232).

74. Yang X., Makaroff C.A., Ma H. 2003 The Arabidopsis MALE MEIOCYTE DEATH1 gene encodes a PHD-finger protein that is required for male meiosis. Plant Cell 15(6), 1281–1295.

75. Cheng C.H., Lo Y.H., Liang S.S., Ti S.C., Lin F.M., Yeh C.H., Huang H.Y., Wang T.F. 2006 SUMO modifications control assembly of synaptonemal complex and polycomplex in meiosis of Saccharomyces cerevisiae. Genes Dev 20(15), 2067–2081. (doi:10.1101/gad.1430406).

76. Maciver S.K., Koutsogiannis Z., de Obeso Fernandez Del Valle A. 2019 ‘Meiotic genes’ are constitutively expressed in an asexual amoeba and are not necessarily involved in sexual reproduction. Biol Lett 15(3), 20180871. (doi:10.1098/rsbl.2018.0871).

77. Friz C.T. 1968 The biochemical composition of the free-living Amoebae *Chaos chaos*, *Amoeba dubia* and *Amoeba proteus*. Comp Biochem Physiol 26, 81–90.

78. Byers T.J. 1986 Molecular biology of DNA in Acanthamoeba, Amoeba, Entamoeba, and Naegleria. Int Rev Cytol 99, 311–341. (doi:10.1016/s0074-7696(08)61430-8).

79. Lohia A. 2003 The cell cycle of *Entamoeba histolytica*. Molecular and Cellular Biochemistry 253(1-2), 217–222.

80. Demin S.Y., Berdieva M.A., Goodkov A.V. 2019 Cyclic Polyploidy in Obligate Agamic Amoebae. Cell and Tissue Biology 13.

81. Maciver S.K. 2016 Asexual Amoebae Escape Muller’s Ratchet through Polyploidy. Trends in parasitology 32(11), 855–862. (doi:10.1016/j.pt.2016.08.006).

82. Spiegel F.W. 2011 Commentary on the chastity of amoebae: re-evaluating evidence for sex in amoeboid organisms. Proceedings Biological sciences / The Royal Society 278(1715), 2096–2097. (doi:10.1098/rspb.2011.0608).

83. Erdos G.W., Raper K.B., Vogen L.K. 1973 Mating Types And Macrocyst Formation In Dictyostelium-Discoideum. Proceedings Of The National Academy Of Sciences Of The United States Of America 70(6), 1828–1830.

84. Erdos G.W., Raper K.B., Vogen L.K. 1976 Effects of light and temperature on macrocyst formation in paired mating types of Dictyostelium discoideum. Journal of bacteriology 128(1), 495–497.

85. Adler P.N., Holt C.E. 1975 Mating type and the differentiated state in Physarum polycephalum. Dev Biol 43(2), 240–253. (doi:10.1016/0012-1606(75)90024-x).

